# Temporal Modulations Reveal Distinct Rhythmic Properties of Speech and Music

**DOI:** 10.1101/059683

**Authors:** Nai Ding, Aniruddh D. Patel, Lin Chen, Henry Butler, Cheng Luo, David Poeppel

## Abstract

Speech and music have structured rhythms, but these rhythms are rarely compared empirically. This study, based on large corpora, quantitatively characterizes and compares a major acoustic correlate of spoken and musical rhythms, the slow (0.25-32 Hz) temporal modulations in sound intensity. We show that the speech modulation spectrum is highly consistent cross 9 languages (including languages with typologically different rhythmic characteristics, such as English, French, and Mandarin Chinese). A different, but similarly consistent modulation spectrum is observed for Western classical music played by 6 different instruments. Western music, including classical music played by single instruments, symphonic, jazz, and rock music, contains more energy than speech in the low modulation frequency range below 4 Hz. The temporal modulations of speech and music show broad but well-separated peaks around 5 and 2 Hz, respectively. These differences in temporal modulations alone, without any spectral details, can discriminate speech and music with high accuracy. Speech and music therefore show distinct and reliable statistical regularities in their temporal modulations that likely facilitate their perceptual analysis and its neural foundations.

## Significance Statement

Speech and music are both rhythmic. This study quantifies and compares the acoustic rhythms of speech and music. A large corpus analysis is applied to speech across languages and to music across genres, including both classical music played by single instruments and ensemble music such as symphonic music, rock music and jazz. The analysis reveals consistent rhythmic properties within the category of speech and within the category of music, but clear distinctions between the two, highlighting potentially universal differences between these fundamental domains of auditory experience.

## Introduction

Rhythmic structure is a fundamental feature of both speech and music. Both domains involve sequences of events (such as syllables, notes, or drum sounds) which have systematic patterns of timing, accent, and grouping (1). A primary acoustic correlate of perceived rhythm is the slow temporal modulation structure of sound, i.e. how sound intensity fluctuates over time (Fig. 1). For speech, temporal modulations below 16 Hz are related to the syllabic rhythm (2) and underpin speech intelligibility (3–5). For music, slow temporal modulations are related to the onsets and offsets of notes (or runs of notes in quick succession), which support perceptual phenomena such as beat, meter, and grouping (1, 6–12). Recently, a number of studies have investigated the neural activation patterns associated with the temporal modulations in the human brain and assessed their relevance to speech and music perception (13–20).

Both speech and music have diverse rhythmic patterns. For example, speech rhythm differs between languages which have sometimes been classified into distinct rhythm categories such as ‘stress-timed’, ‘syllable-timed’, and ‘mora-timed’ (although there is debate whether rhythmic differences between languages are continuous rather than categorical (21)). Furthermore, the details of speech rhythm vary across speakers: people speak at different rates and pause with different patterns. The rhythms of music are even more diverse, with patterns of tempo, grouping, and metrical structure varying dramatically across genres and performances. Yet, despite high diversity, speech and music are both coherent perceptual categories, suggesting features within each domain that are widespread and principled. We explore whether acoustic rhythms constitute one such feature that can separate speech and music into two internally coherent categories.

In speech, temporal modulations of sound intensity show resonance around 4-5 Hz (2, 22, 23), and this acoustic modulation is a correlate of linguistically defined syllabic rhythm (2, 23, 24). Previous studies on speech modulation spectra, however, focused on a small number of speech samples and did not compare the modulation spectra across many languages and large corpora. In music, the resonance properties of the temporal modulations are not yet well characterized, and it remains unclear what units of musical structure underlie modulation peaks, e.g., notes, beats, or higher level metrical patterns.

Here, we first characterize the modulation spectrum of speech and investigate whether the modulation spectrum differs across 9 languages, including American and British English, French, and Mandarin Chinese. The cross-language analysis is designed to reveal whether the speech modulation spectrum is language-specific or whether it highlights a common feature across languages. The languages we analyzed fall into different rhythmic categories. For example, English is typically classified as stress-timed but French as syllable-timed (21). We examine whether the modulation spectrum reflects the rhythmic category of each language or if, rather, it reflects cross-linguistic features constrained by biomechanical and neural properties of the human speech production and perception system (25). Figure 1A exemplifies the materials.

We next characterize the modulation spectra of Western classical tonal-harmonic music, selecting pieces played by single instruments. We investigate whether single-instrument Western classical music, as a coherent perceptual category, shows a consistent modulation spectrum or whether the modulation spectra depend on the instruments being played. In addition, the modulation spectra is computed for Western ensemble music, including symphonic, jazz, and rock music, to test if there is consistency in the modulation spectra across these distinct genres. Figure 1B,C shows examples of the musical materials used. Finally, we quantify the differences between speech and music modulation spectra and test if such differences are perceptually salient.

**Figure 1.**
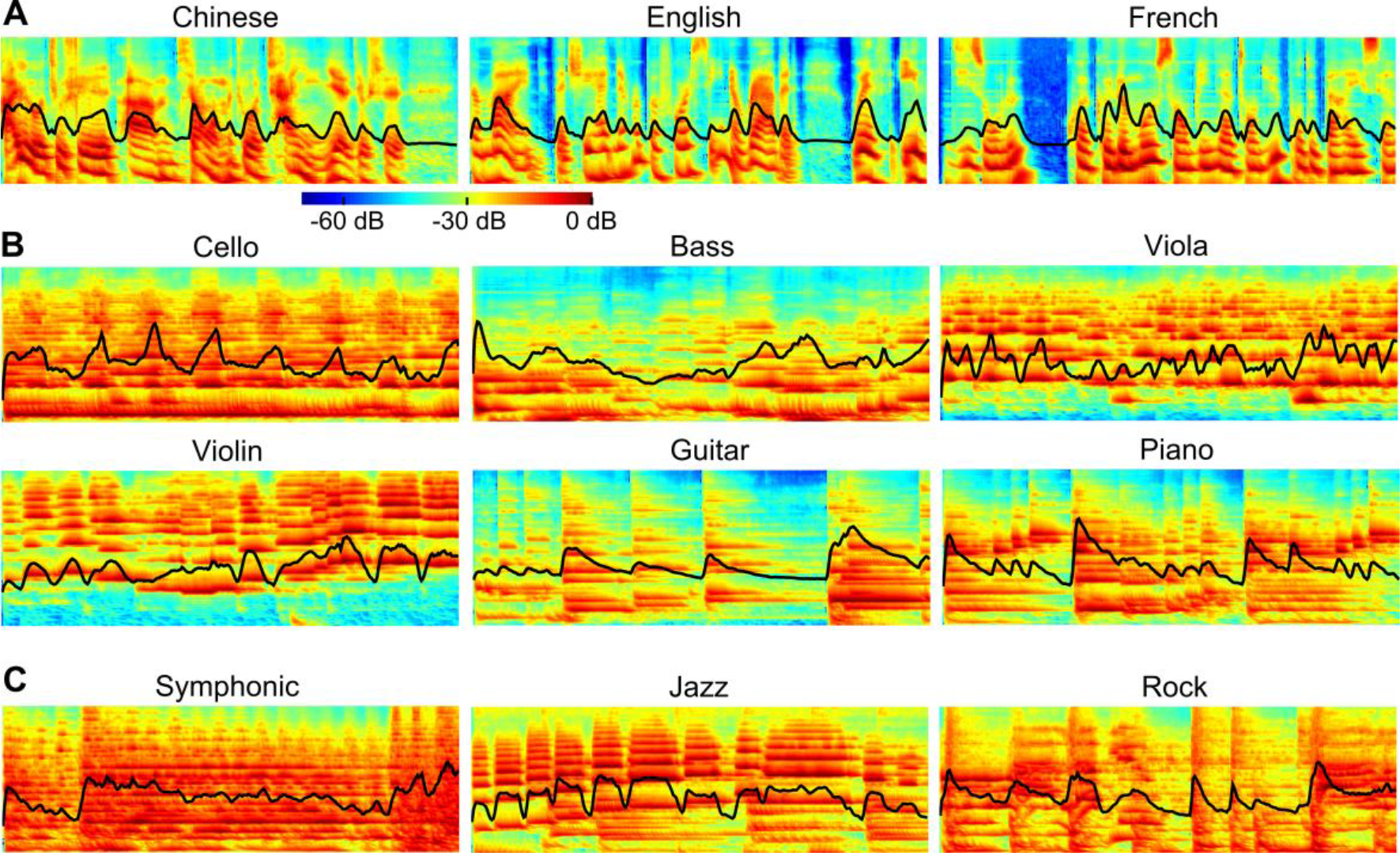
Spectrograms of 3-second randomly chosen excerpts of speech (A), singleinstrument music (B), and ensemble music (C). The spectrograms are simulated using a cochlear model (26). The x-axis denotes time (3 seconds) and the y-axis denotes frequency (180 Hz to 7.24 kHz, on a logarithmic scale). The amplitude of the spectrogram is represented in a logarithmic scale and the maximal amplitude is normalized to 0 dB. The spectrogram summed over frequencies, which reflects how sound intensity fluctuates over time, is superimposed as black lines.

**Figure 2.**
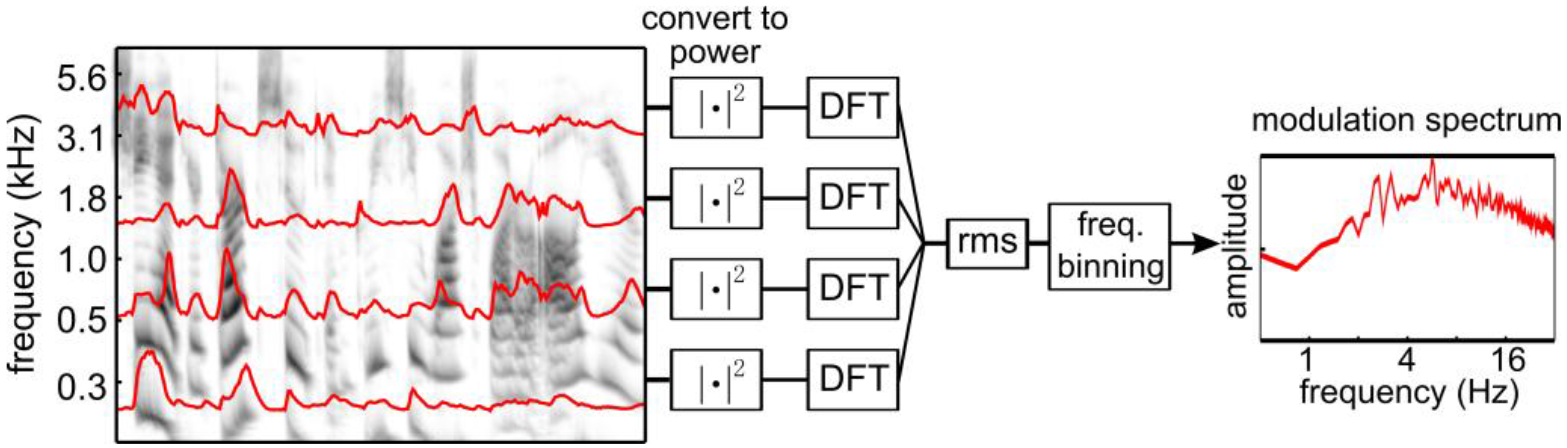
Schematic illustration of how the modulation spectrum is calculated. The sound signal is first decomposed into narrow frequency bands using a cochlear model, and the temporal envelope is extracted in each band (26). Envelopes for 4 frequency bands are illustrated by curves superimposed on the gray-scale figure. The root-mean-square (rms) of the Discrete Fourier Transform (DFT) of all narrowband power envelopes is the modulation spectrum.

## Results

### Speech Modulation Spectrum

The modulation spectrum of speech is extracted using the procedure described in Fig. 2. The discourse-level speech recordings were cut into 6-second duration frames and the modulation spectrum averaged over all frames. The spectrum shows a peak between 4 and 5 Hz, and is highly consistent for the 9 languages we analyzed (Fig. 3A). Additionally, the modulation spectrum is shown to be highly consistent across isolated sentences, audiobooks, and conversational speech in English (Fig. 3B). The averaged peak frequency of the speech modulation spectra is between 4.3 and 5.4 Hz for all tested speech materials (Fig. 3CD; see Fig. S1 for detailed statistics about Fig. 3C; no statistically significant differences are revealed between conditions in Fig. 3D).

**Figure 3.**
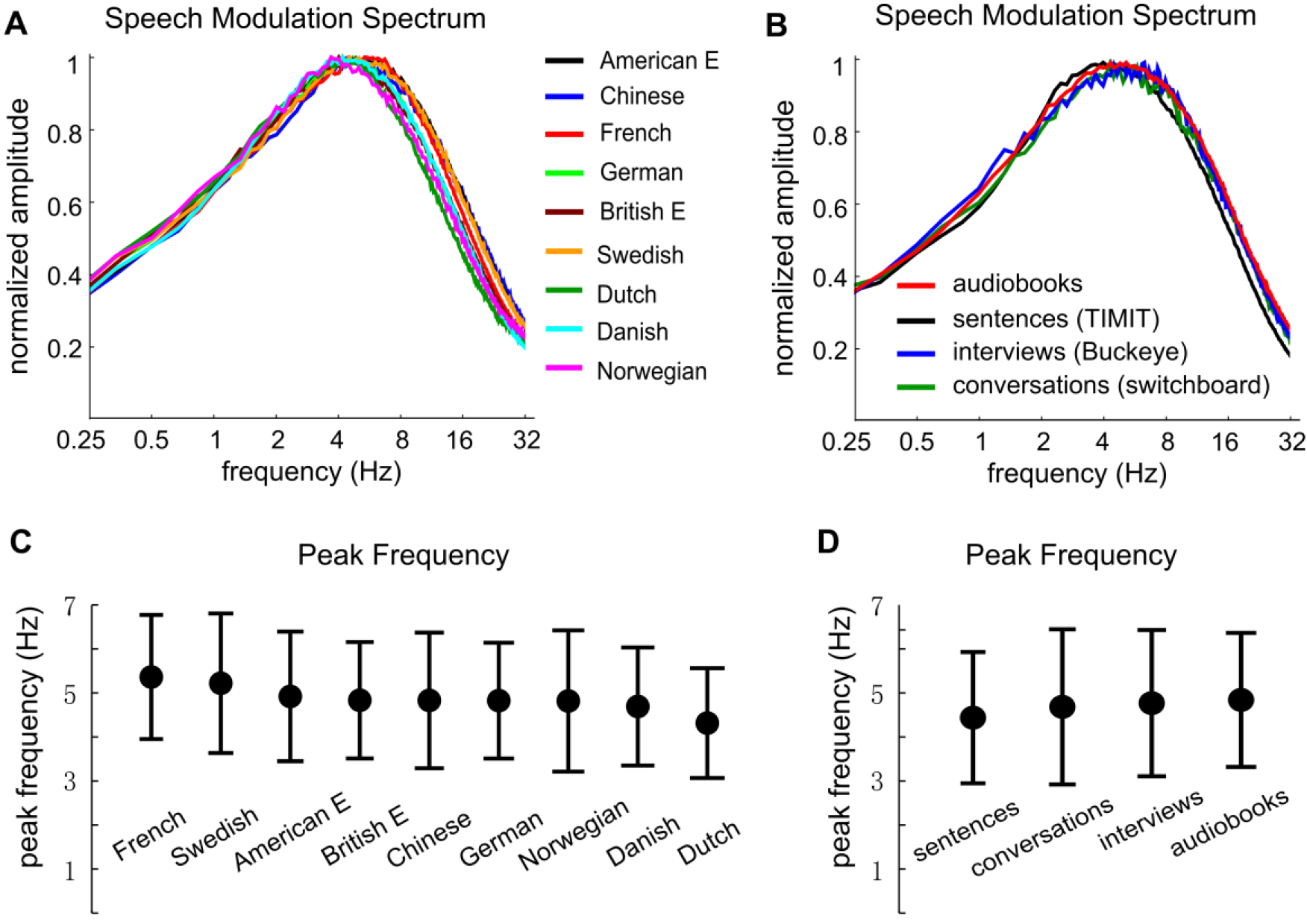
The modulation spectrum of speech. A) The modulation spectrum for naturalistic, discourse-level speech across 9 languages. The modulation spectrum is consistent across languages and shows a peak near 4 Hz. Each spectrum is normalized by its peak amplitude. B) The modulation spectrum for four different corpora of American English. The modulation spectrum is consistent for speech produced in different contexts, including spontaneous and read speech. The peak frequencies for the modulation spectra are shown in C and D. The error bar represents one standard deviation on each side across all the 6-second duration speech recording segments tested for each type of material.

### Music Modulation Spectrum

The modulation spectrum of music shows important differences from that of speech. The modulation spectra of classical music played by four single-voice string instruments (violin, viola, cello, and bass, which typically play one note at a time) are shown in Fig. 4A. Each shows a broad peak between 1 and 2 Hz, substantially lower than the 4-5 Hz peak frequency for speech. The modulation spectra for two multi-voice instruments (piano and guitar), which typically play more than one note at a time, are shown in Fig.4B and do not differ substantially from the average modulation spectrum for the single-voice instruments. Across the six different solo instruments studied here, the modulation spectrum is largely independent of instrument at values below 8 Hz. The modulation spectra for viola and guitar show secondary peaks between 16 and 32 Hz, which may be related to vibratory properties of these instruments.

The modulation spectra of three types of ensemble music are shown in Fig. 4C. The average modulation spectrum of single-voice instruments is also shown for comparison. The modulation spectrum is generally consistent for all analyzed musical styles below 4 Hz, although symphonic music tends to contain more modulation energy at very low frequencies, below about 1.5 Hz. Above 4 Hz the modulation spectra of rock and symphonic music contain more high-frequency energy than the modulation for single-voice instruments and jazz music (the latter two have very similar modulation spectra). For the musical recordings analyzed here, only rock music contains vocals, and therefore it is compared directly with speech in Fig. 4D. Rock music has a broad modulation peak around 2-3 Hz, with considerably more modulation energy than speech in lower frequencies and somewhat less modulation energy than speech between 2 and 16 Hz.

### Differences between Speech and Music Modulation Spectra

The modulation spectrum is directly compared for different languages and musical instruments/genres in Fig. 4D. The spectra show a striking difference: the modulation peak for music is below 2 Hz while the peak for speech is above 4 Hz. Figure 4E shows that the modulation peak for speech (averaged over all materials) is consistently above the peak for music for all musical styles analyzed in the current study. The peak frequency is significantly higher for speech than each of the analyzed music material (*P* < 10^−10^, unpaired t-test between speech and each music category, FDR-corrected).

The pairwise correlation coefficients between speech and music modulation spectra are illustrated in Fig. 4F. The correlation matrix analysis reveals two clusters, one corresponding to speech and the other corresponding to music. The averaged pairwise correlation coefficient is 0.97 ±0.025 (mean ± SD) across languages and 0.98 ±0.016 across musical instruments/genres. In contrast, the mean pairwise correlation between speech and music modulation spectra is 0.51 ±0.13, significantly lower than the pairwise correlation within the speech or music category (*P* < 0.005, unpaired t-test).

**Figure 4.**
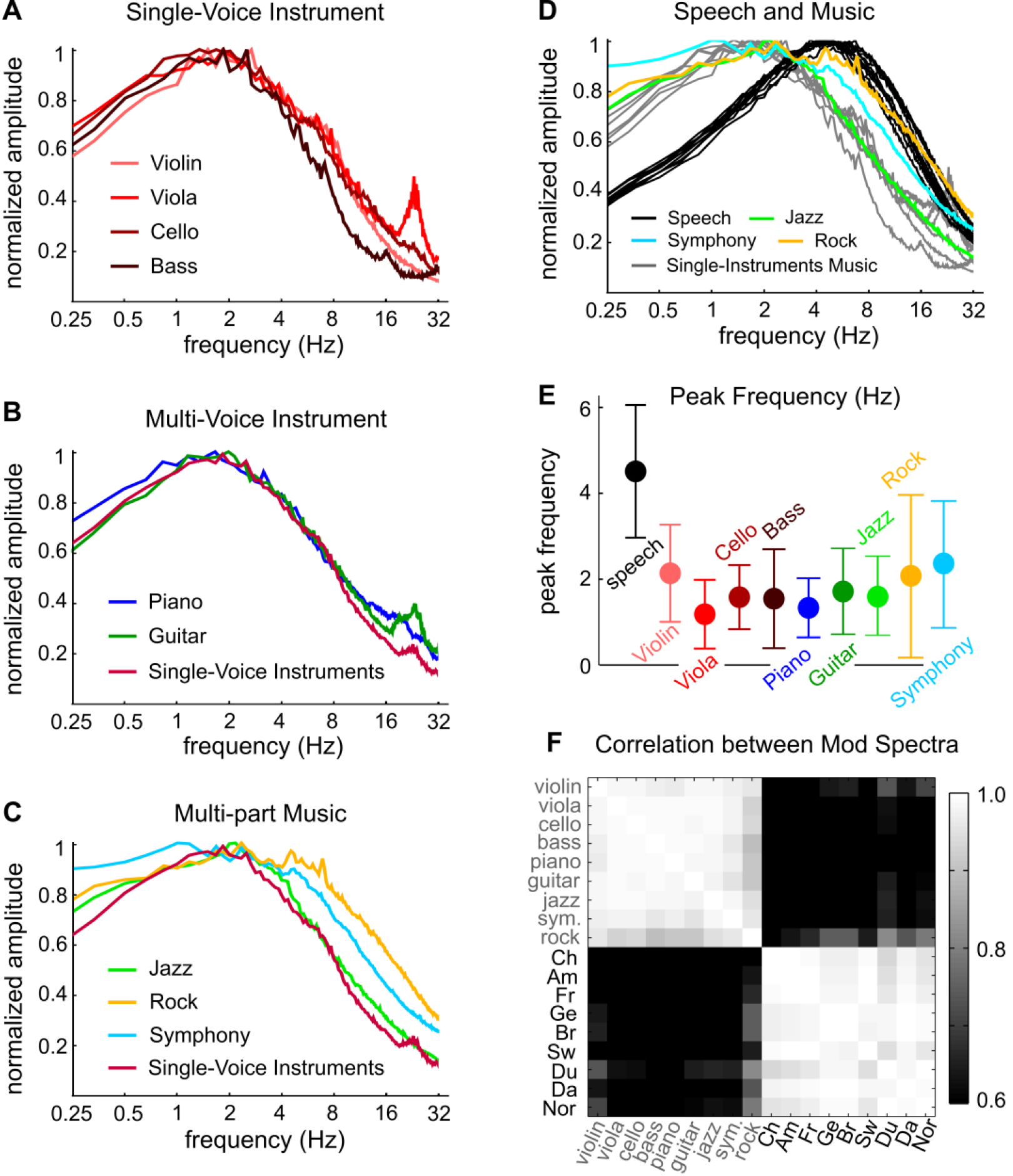
Modulation spectra of Western music. A) The modulation spectrum of classical music played by single-voice string instruments consistently shows a peak below 2 Hz. B) The modulation spectrum of multi-voice instruments, e.g., piano and guitar, is consistent with that of the single-voice string instruments. C) The modulation spectrum of multi-part music, e.g. symphonic music, jazz, and rock, shows a broader peak than single-instrument music, especially for symphonic music and rock. D) The modulation spectrum of speech (reproduced from Fig. 3A), single-instrument music (reproduced from Fig. 4AB), and multi-part music. Speech and music modulation spectra show distinct peak frequencies, and music contains more power at modulation frequencies below 4 Hz. E) The peak frequency of the modulation spectrum is consistently lower for music than speech. The error bar represents 1 standard deviation over all the 6-second duration musical recording segments analyzed for each type of material. F) Pairwise correlation coefficients between speech and music modulation spectra shown in Fig. 4D. See Table S1 and Table S2 for musical materials used in the analysis.

To further quantify the differences between speech and music temporal modulations, we employed a linear classifier to discriminate all 6-s duration speech and music segments. The classifier was trained based on speech and single-instrument music and achieved high classification accuracy (90% and 99% hit rate for speech and music, respectively; 10-fold cross-validation). This classifier was then applied to ensemble music and achieved 100%, 99%, and 88% hit rates for symphonic music, jazz, and rock music, even though these musical genres were not used to train the classifier, suggesting generalizable universal temporal patterning for music in contrast to speech.

To test whether these quantitative differences between speech and music temporal modulations are perceptually salient, listeners were trained to discriminate noise-vocoded speech and noise-vocoded music. The 1-channel noise vocoded stimuli were created to preserve the temporal modulations of speech and music while discarding spectral information, which was achieved by modulating a white noise carrier with the broadband temporal envelope of speech or music.

In the experiment, listeners (n=12) had to discriminate samples of vocoded speech/music with visual feedback (see Procedures). Listeners were instructed that the two categories of sounds were made by two different ‘aliens’; and were not aware that they were vocoded speech and music. The listeners discriminated vocoded speech and music with high accuracy, and the hit rates were 0.95 ±0.04 (mean ± SD over listeners) for both vocoded speech and music. Furthermore, the listeners learned the task quickly and showed no clear learning effects (Fig. S2) These behavioral findings demonstrate that the differences between speech and music temporal modulations are perceptually salient and that listeners can detect this difference with little training.

### Correlates of the Modulation Resonance in Musical Theory

For the music modulation analysis, an important question is whether the spectral peak around 1-3 Hz is related to metrical structure as defined in music theory. A modulation rate of 1-3 Hz is in the range of typical musical beats (1). To investigate how the modulation spectrum might be related to the rate of musical beats for individual musical pieces, we conducted a further analysis based on selecting 25 additional excerpts of music with a clear beat (See Appendix for list). These excerpts were chosen from musical genres in which we could identify extended passages with a strong and steady beat: rock, funk, blues, and electronic music.

As shown in Fig. 5 and quantified in Table S3, in these 25 excerpts the modulation spectrum generally shows a broad peak centered around 1 and 2 times the tempo, with additional narrow peaks at 1, 2, and 4 times the tempo. This suggests that the modulation spectrum peak in music reflects acoustic fluctuations near the beat rate, with additional resonance reflecting a metrical subdivision of the beat, either one metrical level below the beat (2x the beat) or two metrical levels below the beat (4x the beat). Regular subdivision of the beat is a prominent feature of Western music, and research on musical meter suggests that having one or two levels of subdivision below the primary beat level is typical for Western music (7). The temporal modulations provide an acoustic correlate of this musical phenomenon.

**Figure 5.**
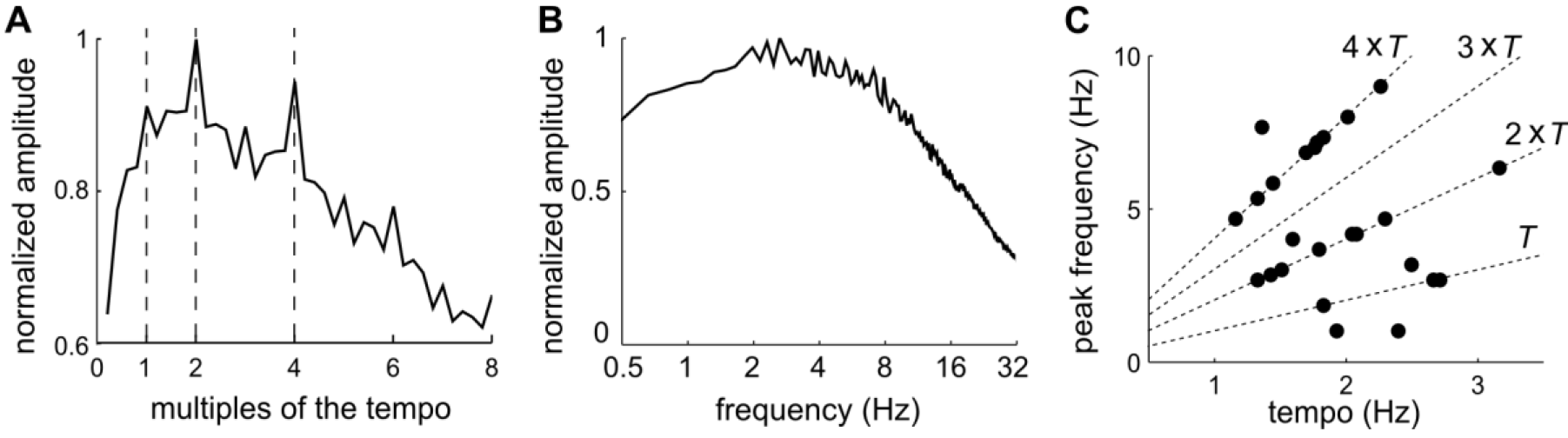
Relationship between the modulation spectra and musical tempi for 25 musical excerpts with a steady beat. A) The modulation spectrum averaged over 25 excerpts. The modulation frequency axis is normalized based on the tempo of each excerpt. The average modulation spectrum shows peaks at the tempo rate and its multiples. The strongest peaks are seen at 2 times the tempo and 4 times the tempo. B) The modulation spectrum when the frequency axis is represented in Hz. C) The relationship between musical tempo and the frequency at which the modulation spectrum shows maximal power, for all 25 excerpts. If the music tempo is denoted as *T* and the modulation spectrum peak frequency is denoted as F, dotted lines are plotted for the relationships: *F = T*, *F* = *2T*, *F* = *3T*, and *F* = *4T*. For most excerpts, the modulation spectrum peak frequency appears at *2T* or *4T*.

## Discussion

This study characterizes in a quantitative manner the slow temporal modulations (< 32 Hz) in sound amplitude for speech and music. The analysis is motivated by the fact that a growing literature in perception and neuroscience research points to the relevance of these modulations in speech and music processing (1, 15, 23). The modulation spectrum of speech shows the greatest power between 2 and 10 Hz, peaking at around 5 Hz, a highly consistent pattern across the 9 languages tested in this study. These results are reminiscent of recent studies showing that the mean syllabic rate of speech is 5-8 Hz across languages (24), indicating universal rhythmic properties of human speech. The 2-10 Hz rhythm is prevalent in the speech communication chain, observed in motor cortex (27) and articulator movements (25, 28) during speech production and in widely distributed cortical areas including auditory cortex during speech perception (29, 30). Therefore, the ~5 Hz rhythm is likely an intrinsic attribute of speech, possibly imposed by the biomechanical properties of the human articulators (25) and the underlying neurodynamical properties of the speech production and perception systems.

The modulation spectrum of Western music also shows consistency across genres and musical instruments, revealing strongest activation between 0.5 and 3 Hz. This finding suggests that the different musical instruments studied here, namely violin, viola, cello, bass, piano, and guitar, do not impose strong constraints on the slow temporal modulations of music. For ensemble music, jazz has a modulation spectrum very similar to that of single-instrument music, while symphonic and rock music show broader spectral speaks. For the music analyzed here, only rock music contains vocals, which may be one reason that its modulation spectrum is closer to the modulation spectrum of speech. The plateau of the musical modulation spectrum, i.e. 0.5-3 Hz, corresponds well to the typical frequency range of musical beats. For example, dance music pieces tend to have tempi between 94 and 176 beats per minute (BPM), which corresponds to a rate of 1.6 Hz to 2.9 Hz for beats (31). Indeed, a subsidiary analysis demonstrates peaks in the music modulation spectra at metrical subdivision values of the basic beat rate or tempo and are thus related to the metrical structure of music.

Although distinct peak modulation frequencies are revealed for speech and music (Fig. 1–2), both speech and music modulation spectra will assume a low-pass 1/f shape when the frequency axis is expressed in a linear rather than logarithmic scale (10, 32, 33). The general 1/f power fall off in the modulation spectra is not unique to speech or music but shared by many other natural sounds (33–35). In a logarithmic frequency scale, the general 1/f trend is removed and a broad peak appears in the 3-5 Hz range for speech and the 0.5-3 Hz range for music. In other words, the distinct spectral peaks for speech and music reflect how these two sound categories deviate from a 1/f modulation spectrum. Human listeners are sensitive to such deviations and can learn to accurately discriminate the two sound categories purely based on temporal information with only a couple of training samples.

Temporal modulations play critical roles in speech and music processing. For speech, it has been proposed that the slow temporal modulations serve as acoustic landmarks to trigger an initial coarse analysis of speech features, which is followed by more finegrained phonetic analysis (36). Consistent with this hypothesis, it has been shown that neural activity in auditory cortex is synchronized to the slow temporal modulations of speech (14, 30, 37). Since the slow temporal modulations correspond to the time scale of syllables, syllable-sized acoustic chunks have been proposed as the basic unit for initial speech analysis (23, 38, 39). For music, beats are fundamental features organizing the temporal structure of music. This study shows that the acoustically salient rhythms correspond to the frequency range for common musical beats and subdivisions, suggesting that these rhythms possibly construct initial time scales for musical analysis (40, 41), in parallel with the landmark hypothesis in speech (36). Indeed, during music listening, cortical activity has been shown to be synchronized to the perceived musical beats (13, 42).

In sum, we characterized the slow temporal modulations of speech and music based on large corpora. The speech recordings (over 25 hours total) contain 9 languages and speech produced in different manners, e.g. during reading or telephone conversation. The music recordings (over 39 hours total) contain single-instrument music from six musical instruments, as well as symphonic, jazz, and rock music. Based on these diverse recordings, a high degree of consistency in the modulation spectra is found within the categories of speech and music – but not across them. These findings suggest that the statistical regularities of slow temporal modulations are intrinsic signatures of speech and music. The current findings suggest that the distinctions between speech and music modulation properties may profitably be taken into account when addressing the distinctions between speech and music perceptual analysis.

It will be important to test to what extent the current findings generalize to languages with diverse prosodic characteristics (43) and to music from non-Western cultures (44). Additionally, it is worth bearing in mind that the analysis here concerns only ‘normal’ speech. The modulation spectrum can certainly be affected by speech production pathology (45) or when a talker speaks exceptionally clearly or slowly (46). It will also be interesting to use the methods presented here to study speech and music that are primarily rhythmic in nature, e.g. metrically regular poems and North Indian tabla music (47, 48), as well as music that contains no clear rhythmic beats e.g. Gregorian chant or Chinese Ch’in music (49).

## Materials and Methods

### Speech Materials

#### American English

Individually pronounced sentences were selected from the TIMIT database (50). All sentences (N = 4,620, totaling 462 minutes of recordings) in the training set of the database were used in the analysis. Continuous discourse-level English speech materials were selected from audiobooks and two speech corpora, i.e. the Buckeye and Switchboard corpus. Four public domain audiobook chapters (from http://librivox.org) were used, read by 4 different talkers (2 female). The duration of each audiobook recording varies between 15 and 83 minutes, and the total duration of the recordings was 140 minutes. The Buckeye speech corpus (51) contains conversations between a talker and an interviewer about everyday topics. Laughter, noise, and speech from the interviewer were removed from the recording based on the labels provided by the corpus. The first 4 audio files from the first 10 talkers in the corpus were used in the analysis, and the total duration of these recordings was 109 minutes. The Switchboard speech corpus contained conversational speech recorded over the telephone and 118 minutes of recordings were analyzed here (23, 52).

#### Mandarin Chinese

Speech recordings in Mandarin Chinese were selected from a radio broadcast, a narrated story, and interviews. A 7-minute radio broadcast recording was selected from the program Sounds of China, on December 1 2013 (http://aod.cnr.cn/). The recording contained a single female talker with no audible background sound. A 6-minute duration narrated story, Cong Cong by Ziqing Zhu, read by a female talker (3 minutes in duration), was also included in the analysis (http://librivox.org/multilingual-short-story-collection-001/). Finally, ten short speeches on Chinese culture were included (http://www.laits.utexas.edu/orkelm/chinese/). The 10 talkers (3 female) were from 10 different provinces in China. The total duration of the recordings was 17 minutes (1-2 minutes for each talker).

#### French

Audiobook recordings in French were selected from 3 chapters of Les Miserables (http://librivox.org/les-miserables-tome-2-by-victor-hugo-1008/), and Le Dernier Jour d’un Condamne (http://archive.org/details/VictorHugo). The 4 sections were read by 4 different talkers (2 female). The total duration of the recordings was 64 minutes and the duration of each book chapter varies between 12 and 21 minutes. Additionally, the French materials also include short passages from the EUROM corpus, which is detailed below.

#### Seven European Languages

Materials include all the short passages from the EUROM corpus, which are in 7 different languages, i.e. Danish, Dutch, English, French, German, Norwegian, and Swedish. The corpus includes 60-64 speakers for each language. Three to five passages were recorded for each speaker and additional 15-20 passages were recorded from 10 of the speakers for each language. The total duration of recordings ranges from 44 minutes to 130 minutes across the 7 languages.

All speech recordings were resampled to 16 kHz. All recordings not from a standard speech corpus were judged as natural and representative of American English, French, or Mandarin English by a native speaker of the language. Figure 1A shows the auditory spectrogram simulated using a cochlear model (26) for a 3 second excerpt of the audiobook materials from each of the three languages.

### Musical Materials

#### Single instrument materials

Single instrument music was analyzed for six instruments: violin, viola, cello, bass, guitar, and piano. Table S1 shows the different instruments analyzed, the duration of material for each instrument, representative composers whose work was included in the analysis, and one example piece for each instrument (for a full catalog of musical pieces included in this study, please see Appendix). Figure 1B shows the auditory spectrogram simulated using a cochlear model for a 3 second excerpt of music from each of the instruments analyzed.

#### Ensemble recordings

Table S2 shows the different genres analyzed, duration of material for each genre, composers whose work was included in the analysis, and one example piece for each genre. (Of the three genres analyzed, symphonic and jazz were purely instrumental music, while rock was instrumental plus vocals.) Figure 1C shows the auditory spectrogram simulated using a cochlear model for a 3 second excerpt for each of the ensemble musical categories analyzed.

#### Recordings for tempo analysis

Twenty-five excerpts of music with a clear beat were chosen, which span a wide range of tempi (55 – 190 BPM) and ranged in duration from 60 s to 286 seconds (average duration = 163 s). Most excerpts combined instruments and vocals (five were purely instrumental). These excerpts were selected by co-author HB, an amateur drummer (11 years experience), who verified that each except had a steady tempo and who determined the tempo of each excerpt by drumming along with it.

### Temporal Envelopes

The temporal envelope was extracted using a cochlear model (26). The model filters the sound signal into 128 frequency bands, evenly distributed over a logarithmic frequency scale between 180 Hz and 7 kHz (Fig. 2). In each band, in order to extract the temporal envelope, the filtered sound signal is half-wave rectified, smoothed by a single-pole low-pass filter with a time constant of 8 ms, squared, and then decimated to 100 Hz (22).

### Modulation Spectrum

A Discrete Fourier Transform (DFT) was applied to each narrowband power envelope. Before the DFT, the sentence-level recording was cut or zero padded to 6 seconds in duration and longer speech/music recordings were cut into 6-second duration samples. No smoothing window was applied. The modulation spectrum is the absolute value of the Fourier coefficients weighted by the modulation frequency. If the Fourier coefficient is denoted as *s*(*f*), the modulation spectrum is *f*^1/2^|*s*(*f*)|. The modulation spectrum created in this way is equivalent to filtering the temporal envelope using a logarithmically spaced filter bank and calculating the root mean square value (RMS) of the output of each filter, which is the traditional way of calculating the modulation spectrum (22, 53). The final modulation spectrum is the RMS over the carrier frequency bands and is normalized by its maximal value between 0.25 and 32 Hz. The modulation spectrum is finally averaged within each corpus by calculating the RMS value.

### Linear Classification of Speech and Music Modulation Spectrum

A linear classifier was employed to classify the modulation spectrum of individual 6-sec duration speech and music samples. In this analysis, it was assumed that the modulation spectrum for each sound category, i.e., speech or music, was subject to a multidimensional Guassian distribution. Parameters of each Guassian distribution were fitted based on training samples. Based on the fitted Gaussian distribution, a linear hyperplane was used to separate speech samples and music samples. In the classification analysis and the correlation analysis in Fig. 2F, the modulation spectrum was binned and the bin size was 0.5 octave.

### Behavioral Classification of Vocoded Speech and Music

To test if listeners are sensitive to the differences between speech and music modulation spectra, we asked them to discriminate 1-channel noise vocoded speech and music samples. Materials include 7 European languages and 9 musical instruments/genres. Each sample was 4-seconds in duration and was randomly chosen from the relevant corpora. The temporal envelope of each speech/music sample (low-pass filtered at 4 kHz) was extracted using the Hilbert transform and low-pass filtered below 32 Hz. The temporal envelope was then used to modulate a band-limited white noise (125 Hz – 4 kHz) to generate the 1-channel vocoded speech/music.

Twelve listeners (21-24 yrs old, mean 22 yrs old; 7 male) participated in the experiment. All subjects reported normal hearing and no formal musical training. Written informed consent was obtained from each participant prior to the experiment. In the experiment, the listeners sat in a quiet room before a screen and sounds were delivered using insert earphones (Sennheiser CX213). They were instructed to discriminate the sounds made by two aliens, without knowing that the two sound categories were in fact vocoded speech and music.

The experiment was divided into 3 stages. In stage 1, the listeners clicked on each alien to listen to one 4-s duration sample sound. In stage 2, the listeners listened to 10 vocoded stimuli and had to judge which alien made the sound by pressing different buttons. Visual feedback, i.e. right or wrong, was given after each response. Half of the 10 stimuli were vocoded speech and the half were vocoded music. Each sample was from a different language or musical instrument/genre that was randomly selected online for each listeners. If the averaged correct rate in stage 2 is above 90%, the listeners were allowed to continue to the third stage, in which they had to discriminate 126 vocoded stimuli with visual feedback. Half of the stimuli (*N* = 63) were vocoded speech (9 samples for each of the 7 languages) and the other half were vocoded music (7 samples for each of the musical instrument/genre). All listeners successfully proceeded to stage 3 after stage 2. The reported hit rates were based on stage 3, while the results from stage 2 were shown in Fig. S2.

Since speech tends to contain pauses while music does not, to ensure that pauses are not the only cue used to discriminate vocoded speech and music, we further created a condition in which pauses longer than 250 ms were removed. For this conditoin, we first randomly selected 6-seconds duration segments from speech and music corpora with the criterion that the total duration of long pauses was shorter than 1.5 seconds. Long pauses were then removed from the noise-vocoded stimulus, after which the first 4 seconds of the stimulus was retained for the experiment. Six listeners participated in this no-pause condition and the other six listeners participated in the condition in which the pauses in speech were not manipulated. The hit rates from the no-pause condition were not statistically different from the original condition in which pauses were not removed (*P* > 0.1. unpaired t-test), and therefore results from these two conditions were pooled together.

## Acknowledgements

We thank Erik Broess of Tufts University for help in selecting the music materials and Jess Rowland for help in creating the music catalog. Work supported by NIH 2R01DC05660 (DP), National Natural Science Foundation of China 31500873 (ND), Fundamental Research Funds for the Central Universities (ND), and Zhejiang Provincial Natural Science Foundation of China LR16C090002 (ND).

## Supporting Figures

**Figure S1.**
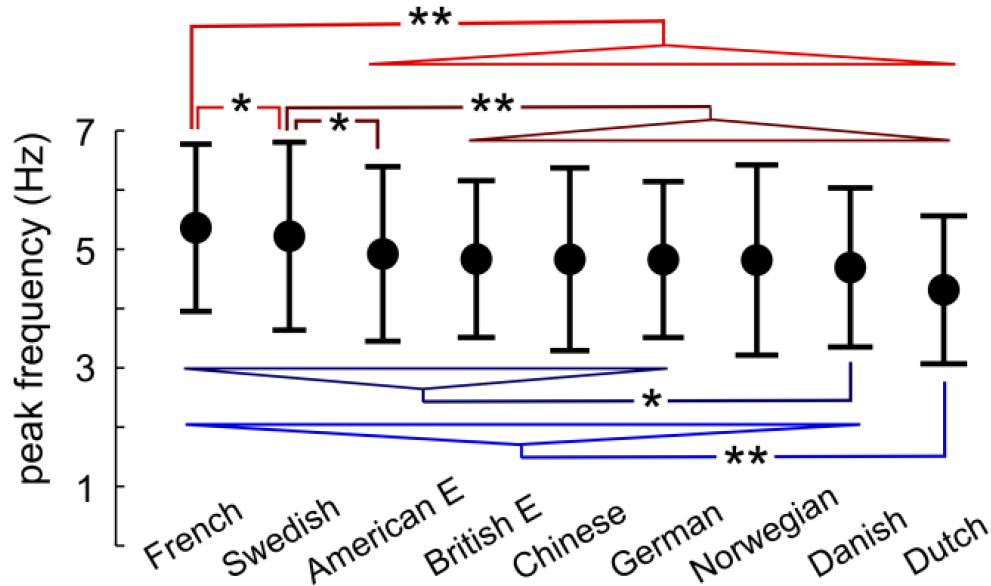
Statistical analysis of the differences between modulation peak frequencies across languages. Statistically significant differences are marked by stars (\**P* < 0.05 and * *P* < 0.05, FDR corrected). Only 4 languages, i.e., French, Swedish, Danish, and, Dutch, show significant differences from other languages, and comparisons based on each of the
4 languages is shown by a different color.

**Figure S2.**
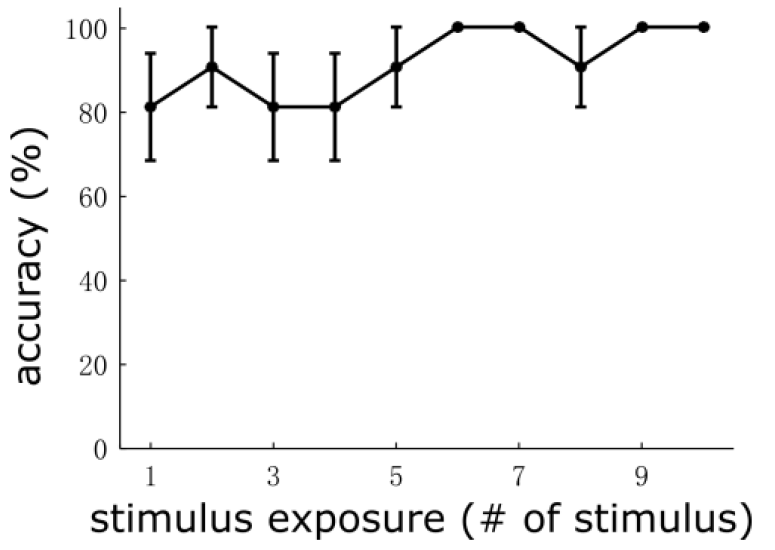
Behavioral discrimination of vocoded speech and music. Data are based on stage 2 of the behavioral experiment (see Procedures for details). In brief, in stage 1, the listeners were exposed to one sample of vocoded speech and one sample of vocoded music (4 seconds in duration). In stage 2, the listeners had to discriminate 10 novel vocoded stimuli, with visual feedback. Even for the very first stimulus in stage 2, they achieved greater than 80% accuracy, indicating that the listeners can capture the differences between vocoded speech and music based on even one example per category. Error bars show 1 standard error on each side.

**Table S1.** Solo instrumental music included in this study.

| Instrument | Representative Composers | Duration (minutes) | # of artists | Example Piece |
| --- | --- | --- | --- | --- |
| Piano | Beethoven, Mozart | 230 | 4 | Beethoven, Sonata Op. 27 No. 2 “Moonlight” in C Sharp |
| Cello | Bach, Ysaÿ | 139 | 2 | Bach, J.S. No. 1, S. BWV 1007, G major |
| Bass | Bach | 60 | 1 | Bach, J.S. Suites, violoncello, BWV 1008 |
| Viola | Bach, Hindemith | 210 | 3 | Bach, J.S. Suite no. 1 in G major BWV 1007 |
| Violin | Bach, Ysaÿ | 204 | 2 | Bach, J.S. Sonata no. 1 in g minor, BWV 1001 |
| Guitar | Bach, Sanz | 216 | 2 | Sanz Pavanas |

**Table S2.**
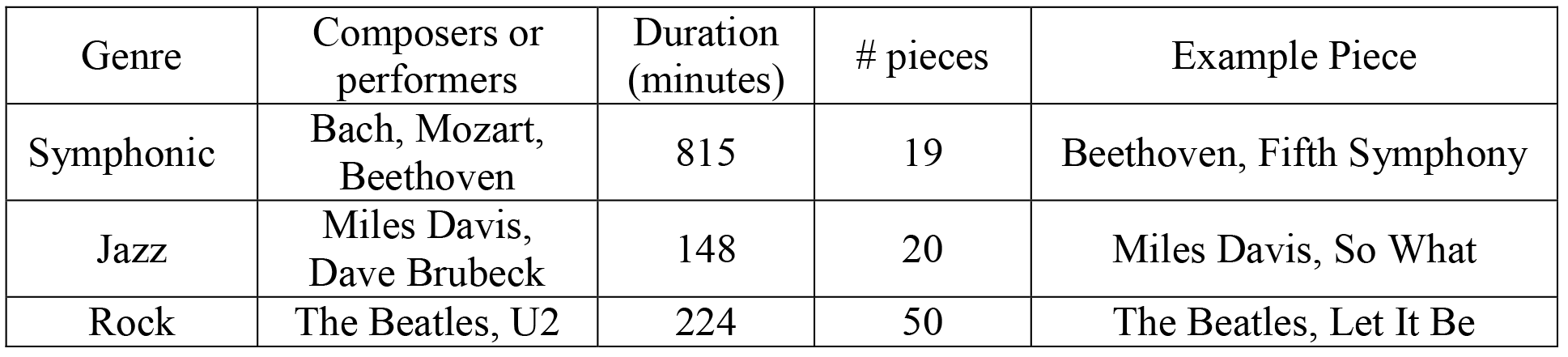
Ensemble music included in this study.

**Table S3.** The relationship between musical tempo and peaks in the music modulation spectrum for the musical excerpts shown in Figure 5.The 2^nd^ row shows whether the power at a frequency of interest (e.g., the tempo rate or its harmonically related frequencies) is significantly larger than the power averaged over neighboring frequency bins (paired t-test based on 25 excerpts). If the tempo is denoted by *T* and the frequency of interest is denoted as f the neighboring bins considered here fall between *f – T* and *f*+ *T*. A significant p-value means that the power at the frequency interest, *f*, is larger than the power of any frequency bin falling between *f – T* and *f* + *T*. The average modulation spectrum shows a statistically significant peak at *T*, 2*T*, 3*T*, and 4*T*. The 3^rd^ row shows the number of songs showing a local maximum at each frequency of interest. A local maximum is caculated between *f – T* and*f* + *T*. Most excerpts show a spectral peak at 2*T* or 4*T*.

| | | Tempo | $2 \times \text{tempo}$ | $3 \times \text{tempo}$ | $4 \times \text{tempo}$ |
| --- | --- | --- | --- | --- | --- |
| Narrowband<br>Envelope | P-value | 0.006 | $<10^{-4}$ | 0.001 | $<10^{-4}$ |
|  | % of songs<br>showing a peak | 40% | 72% | 56% | 68% |

## Appendix <u>Piano</u>

Mozart Piano Sonatas K. 310, K. 331, K. 533/494

Perahia, Murra, pianist.

Sony Classical, SK48233 (1995)

> *Mozart WA, Piano Sonata in A minor, K 330*
>
> *Mozart WA, Piano Sonata in A major, K 331*
>
> *Mozart WA, Piano Sonata in F major, K 533/494*

Beethoven: Sonata, Op.27,No.2 / Franck: Prelude, choral et fugue / Brahms: Paganini

Variations

Kissin, Evgeny, pianist.

RCA, BMG 09026-68910-2 (1998)

> *Beethoven, L. Sonata Op. 27 No. 2 ‘Moonlight’ in C Sharp*
>
> *Franck, C. Prelude, Choral et Fugue*
>
> *Brahms, J. Paganini Variations*

Beethoven: Piano Sonatas, Op.81a & 106

Brendel, Alfred, pianist.

Phillips Import, Philips 446 093-2 (1996)

> *Beethoven, L. Sonata for piano No. 29 in B flat major (“Hammerklavier“) Op. 106*
>
> *Beethoven, L. Sonata for piano No. 26 in E flat major (“Les Adieux“) Op. 81a*

Mozart: Klaviersonaten

Pogorelich, Ivo, pianist.

Dg Imports, DG4377632 (1995)

> *Mozart WA, Fantasia for Piano in D minor K 397*
>
> *Mozart WA, Piano Sonata in G major K 283*
>
> *Mozart WA, Piano Sonata in A major K 331*

## Cello

Unaccompanied cello suites. Johann Sebastian Bach.

Ma, Yo-Yo, cellist.

Columbia, M2K 37867 (1983)

> *Bach JS. No. 1, S. BWV 1007, G major*
>
> *Bach JS. No. 4, S. BWV 1010, E-flat major*
>
> *Bach JS. No. 5, S. BWV 1011, C minor*
>
> *Bach JS. No. 2, S. BWV 1008, D minor*
>
> *Bach JS. No. 3, S. BWV 1009, C major*
>
> *Bach JS. No. 6, S. BWV 1012, D major*

Cello Sonatas

Epperson, Gordon, cellist.

Centaur, CRC 2228 (1995)

> *Eugène Ysay. Sonate pour violoncelle seul, op. 28*
>
> *George Crumb. Sonata for solo violoncello*
>
> *Zoltán Kodály. Sonate pour violoncelle seul, op. 8*

## Bass

Unaccompanied cello suites: performed on double bass

Meyer, Edgar, bassist.

Sony Classical, SK 89183 (2000)

> *Bach, JS. Suites, violoncello, BWV1008*
>
> *Bach, JS. Suites, violoncello, BWV 1007*
>
> *Bach, JS. Suites, violoncello, BWV 1011*

## Viola

Bach, JS. Six cello suites, on viola

Callus, Helen, violist.

Analekta, AN 2 9968-9 (2011)

> *Bach, JS. Suite no. 1 in G major BWV 1007*
>
> *Bach, JS. Suite no. 2 in D minor, BWV 1008*
>
> *Bach, JS. Suite no. 3 in C major BWV 1009*
>
> *Bach, JS. Suite no. 4 in Eflat major BWV 1010*
>
> *Bach, JS. Suite no. 5 in C minor BWV 1011*
>
> *Bach, JS. Suite no. 6 in D BWV 1012 (transposed in G major)*

Hindemith, Viola Sonatas

Levin, Robert D. Sonatas for viola alone; Sonatas for viola and piano

Kashkashian, Kim, violist.

ECM New Series, ECM 1330-32 (1988)

> *Paul Hindemith. Sonatas for viola alone. op. 31/4*
>
> *Paul Hindemith. Sonatas for viola alone op. 25/1*
>
> *Paul Hindemith. Sonatas for viola alone op. 11/5*

Watras, Melia. Viola solo

Fleur de Son Classics, FDS 57962 (2004)

> *Arad, Atar. Sonata*
>
> *Bach, Johann Sebastian. Chromatische Fantasie und Fuge. Fantasia*.
>
> *Corigliano, John. Fancy on a Bach air*
>
> *Waggoner, Andrew. Collines parmi étoiles*
>
> *Stravinsky, Igor. Elegy*.
>
> *Prestini, Paola. Sympathique*.
>
> *Penderecki, Krzysztof. Cadenza*

## Violin

Ysaeÿ Eugenè. Six sonatas for solo violin, op. 27

Murray, Tai, violinst.

Harmonia Mundi USA, HMU 907569 (2012)

> *Ysaeÿ Eugène. Sonata no. 1*
>
> *Ysaeÿ Eugène. Sonata no. 2*
>
> *Ysaeÿ Eugène. Sonata no. 3*
>
> *Ysaeÿ Eugène. Sonata no. 4*
>
> *Ysaeÿ Eugène. Sonata no. 5*
>
> *Ysaeÿ Eugène. Sonata no. 6*

Bach, JS. The complete sonatas and partitas for solo violin. Vol. 1

Ross, Jacqueline, violinst.

Gaudeamus, GAU 358 (2007)

> *Bach JS. Sonata no. 1 in g minor, BWV 1001*
>
> *Bach JS. Partita no. 1 in B minor, BWV 1002*
>
> *Bach JS. Sonata no. 2 in A minor, BWV 1003*

Bach, JS. The complete sonatas and partitas for solo violin. Vol. 2.

Ross, Jacqueline.

Gaudeamus, GAU 359 (2007)

> *Bach JS. Partita no. 2 in D minor, BWV 1004*
>
> *Bach JS. Sonata no. 3 in C, BWV 1005*
>
> *Bach JS. Partita no. 3 in E, BWV 1006*.

## Guitar

Segovia, AndrEugenés. Art of Segovia

Deutsche Grammophon, 289 471 697-2 (2002)

> *Handel, George Frideric. Suite for harpsichord no. 4 in D minor, HWV 43 7*.
>
> *Sarabande*
>
> *Bach JS. Suite for violoncello solo no. 1 in G major, BWV 1007. Prélude*
>
> *Bach JS. Partita for violin solo no. 1 in B minor, BWV 1002. Tempo de bourrée*
>
> *Bach JS. Suite for violoncello solo no. 3 in C major, BWV 1009. Courante*
>
> *Bach JS. Partita for violin solo no. 3 E major, BWV 1006.Gavotte en rondeau*
>
> *Aria e corrente*
>
> *Frescobaldi, Girolamo. Sonata in C minor, K. 11 (L. 352)*
>
> *Scarlatti, Domenico. Nouvelles suites de pièces de clavecin*.
>
> *Rameau, Jean-Philippe. Menuet in G major*.
>
> *Manuel Ponce after Nicolò Paganini. Andantino variato*
>
> *Chopin, Frédéric. 24 préludes, op. 28. No. 7 in A major*
>
> *Mendelssohn, Felix. String quartet in E flat major, op. 12. Canzonetta*
>
> *Franck, César. L’Organiste, FWV 41, Sept pièces en mi bémol majeur et mi*
>
> *bémol mineur. Quasi lento, Andantino poco allegretto*
>
> *Mussorgsky, Modest. Pictures at an exhibition. Il vecchio castello*
>
> *Grieg, Edvard. Lyric pieces IV, op. 47. Melodie*
>
> *Debussy, Claude. Préludes, livre 1. La fille aux cheveux de lin*
>
> *Scriabin, Alexander. 5 preludes, op. 16. No. 4 in E flat minor*
>
> *Albéniz, Isaac. Suite española. Asturias. Leyenda-Preludio*
>
> *Albéniz, Isaac. Piezas características. Zambra granadina*
>
> *Segovia, Andrés. Estudio sin luz*
>
> *Rodrigo, Joaquín. Fantasía para un gentilhombre for guitar and small orchestra*.
>
> *Danza de las hachas*
>
> *Jordà, Enrique. Symphony of the Air*

Bream, Julian. Baroque Guitar

RCA Victor Gold Seal, RCA 60494 (1991)

> *Sanz, Gaspar. Pavanas*
>
> *Sanz, Gaspar. Galliardas*
>
> *Sanz, Gaspar. Passacalles*
>
> *Sanz, Gaspar. Canarios*
>
> *Guerau, Francisco. Villano*
>
> *Guerau, Francisco. Canario*
>
> *Bach, JS. Prelude in D minor, BWV 999*
>
> *Bach, JS. Fugue in A minor, BWV 1000*
>
> *Weiss, Sylvius Leopold. Passacaille*
>
> *Weiss, Sylvius Leopold. Fantasie*
>
> *Weiss, Sylvius Leopold. Tombeau sur la mort de M. Comte de Logy*
>
> *de Visée, Robert. Suite in D minor*
>
> *Frescobaldi, Girolamo. Aria con variazione detta la Frescobalda arr. Segovia*
>
> *Scarlatti, Domenico. Sonata in E minor, K. 11*
>
> *Scarlatti, Domenico. Sonata in E minor, K. 87, arr. Bream [K. 11], Segovia [K. 87]*
>
> *Cimarosa, Domenico. Sonata in C# minor*
>
> *Cimarosa, Domenico. Sonata in A, arr. Bream*.

## Symphonies

The Beethoven Symphonies 1-9 Live From the Edinburgh Festival

Mackeras, Sir Charles, conductor.

Hyperion UK, CDS44301/5 (2007)

> *Beethoven L. Symphony #1*
>
> *Beethoven L. Symphony #2*
>
> *Beethoven L. Symphony #3*
>
> *Beethoven L. Symphony #4*
>
> *Beethoven L. Symphony #5*
>
> *Beethoven L. Symphony #6*
>
> *Beethoven L. Symphony #7*
>
> *Beethoven L. Symphony #8*
>
> *Beethoven L. Symphony #9*

Mozart: Symphonies Nos. 39 & 40

Jacobs, René, conductor. Freiburger Barockorchester.

Harmonia Mundi, HMC901959 (2010)

> *Symphony 39 in E-flat major K 543*
>
> *Symphony 40 in G minor K 550*

Mozart: Symphony No. 41; La Clemenza di Tito Overture

Brüggen, Frans, conductor. Orchestra of the Eighteenth Century.

Philips Digital Classics, Phillips 420 241-1 (2010)

> *Mozart WA, Symphony 41*
>
> *Mozart WA, La Clemenza di Tito Overture*

Bach: Brandenburg Concertos

Alessandrini, Rinaldo, conductor. Concerto Italiano.

Naïve, OP 30412 (2005)

> *Bach JS. Brandenburg Concerto No. 1 in F major, BWV 1046*
>
> *Bach JS. Brandenburg Concerto No. 2 in F major, BWV 1047*
>
> *Bach JS. Brandenburg Concerto No. 3 in G major, BWV 1048*
>
> *Bach JS. Brandenburg Concerto No. 4 in G major, BWV 1049*
>
> *Bach JS. Brandenburg Concerto No. 5 in D major, BWV 1050*
>
> *Bach JS. Brandenburg Concerto No. 6 in B flat major, BWV 1051*

## Jazz

Davis, Miles. Kind of Blue (or. 1959)

Columbia/Legacy, CN 90887 (2004)

> *So what*
>
> *Freddie Freeloader*
>
> *Blue in green*
>
> *All blues*
>
> *Flamenco sketches*
>
> *Flamenco sketches (alternate take)*

Dave Brubeck Quartet. Time Out (or. 1959)

Columbia/Legacy, CN 88697 (2009)

> *Blue rondo àla Turk*
>
> *Strange meadow lark*
>
> *Three to get ready*
>
> *Kathy’s waltz*
>
> *Everybody’s jumpin′*
>
> *Pick up sticks*
>
> *The Dave Brubeck Quartet live at Newport. St. Louis Blues*
>
> *Waltz limp*
>
> *Since love had its way*
>
> *Koto song*
>
> *Pennies from heaven*
>
> *You go to my head*
>
> *Blue rondo àla Turk*
>
> *Take five*

## Rock

The Beatles. The Beatles 1967-1970

Apple Records, CDP 0777 7 97039 2 0 (1993)

> *Strawberry Fields Forever*
>
> *Penny Lane*
>
> *Sgt. Pepper’s Lonely Hearts Club Band*
>
> *Here Comes the Sun*
>
> *Come Together*
>
> *Something*
>
> *Octopus’s Garden*
>
> *Let It Be*
>
> *Across the Universe*
>
> *The Long and Winding Road*

The Beatles. The Beatles 1962-1966

Apple Records, CDP 0777 7 97036 2 3 (1993)

> *Love Me Do*
>
> *Please Please Me*
>
> *From Me To You*
>
> *She Loves You*
>
> *I Want To Hold Your Hand*
>
> *All My Loving*
>
> *Can’t Buy Me Love*
>
> *A Hard Day’s Night*
>
> *And I Love Her*
>
> *Eight Days A Week*
>
> *I Feel Fine Ticket To Ride*
>
> *Yesterday*
>
> *Help!*
>
> *You’ve Got To Hide Your Love Away*
>
> *We Can Work It Out*
>
> *Day Tripper*
>
> *Drive My Car*
>
> *Norwegian Wood (This Bird Has Flown)*
>
> *Nowhere Man*
>
> *Michelle*
>
> *In My Life*
>
> *Girl*
>
> *Paperback Writer*
>
> *Eleanor Rigby*
>
> *Yellow Submarine*

U2. The Best of 1980-1990

Island, ADV7963-2 (1998)

> *Pride (In The Name Of Love)*
>
> *New Year’s Day*
>
> *With Or Without You*
>
> *I Still Haven’t Found What I’m Looking For*
>
> *Sunday Bloody Sunday*
>
> *Bad*
>
> *Where The Streets Have No Name*
>
> *I Will Follow*
>
> *Unforgettable Fire*
>
> *Sweetest Thing*
>
> *Desire*
>
> *When Love Comes To Town*
>
> *Angel Of Harlem*
>
> *All I Want Is You*

## 30 Rock Songs for the Tempo Analysis

CD: Deeds, Not Words

Player: Max Roach

CD Info: Riverside Label, B000000YH2, released in 1958

Songs: Deeds, Not Words

CD: Attack and Release

Player: The Black Keys (Dan Auerbach and Patrick Carney)

CD Info: Danger Mouse (producer), Nonesuch (label), B0014QABX0, released in 2008

Songs: All You Ever Wanted

CD: Rudeboy

Player: Zeds Dead

CD Info: San City High Records, Digital Download B004F7A71G, released in 2010

Songs: Rude Boy

CD: John the Conqueror

Player: John the Conqueror (written by Pierre Moore)

CD Info: Alive Natural Sound Records, B009369YMM, released in 2012

Songs: All Alone

CD: Meters

Player: The Meters (Art Neville, Ziggy Modeliste, Leo Nocentelli, George Porter Jr.)

CD Info: Sundazed Music, Inc, B0000365IM, released in 1969

Songs: Ease Back

CD: Electriclarryland

Player: The Butthole Surfers ( Gibby Haynes and Paul Leary)

CD Info: Capitol (Label), B000002TS3, produced by Paul Leary, Steve Thompson, released in 1996

Songs: Pepper

CD: Every Hour Is A Dollar Gone

Player: Patrick Sweany

CD Info: Nine Mile Records, B000QUTS0M, released in 2007

Songs: Them Shoes

CD: Conjure

Player: Voodelic (Earl Lundy)

CD Info: Topisaw Dawg (label), B002PLEFZS, Perry Trest (producer), released in 2009

Songs: Universal Screw

CD: The Great Escape of Leslie Magnafuzz

Player: Radio Moscow (Parker Griggs)

CD Info: Alive Naturalsound (label), B005IY3DH0, Park Griggs (producer), released in 2011

Songs: Creepin’

CD: By A Thread

Player: Gov’t Mule (Warren Haynes)

CD Info: Evil Teen Records (Label), B002MBAJ4M, produced by Gordie Johnson and Warren Haynes, released in 2009

Songs: Steppin’ Lightly

CD: Burnt Offering

Player: The Budos Band

CD Info: Daptone Records, B00N1CHKLQ, released in 2014

Songs: Aphasia

CD: The Glorious Dead

Player: The Heavy (Chris Ellul, Spencer Page, Kelvin Swaby, Daniel Taylor)

CD Info: Counter (Label), B0088MJO9A, Produced by Paul Corkett, released in 2012

Songs: The Big Bad Wolf

CD: Come On In

Player: R.L Burnside

CD Info: Fat Possum Records, B000008UMZ, released in 1998

Songs: Let My Baby Ride

CD: Funkentelechy Vs. The Placebo Syndrome

Player: Parliament (George Clinton)

CD Info: Casablanca (Label), B000001FCK, released in 1977, 1990 (reissue)

Songs: Bop Gun (Endangered Species) and Flashlight

CD: Bacano

Player: Orgone

CD Info: Killion Floor Sound (Label), B001NW4JJW, released in 2008

Songs: Be In Here

CD: Tres Hombres

Player: ZZ Top (Billy Gibbons, Frank Beard, Dusty Hill)

CD Info: Warner Bros. (Label), B000CCD0HQ, released in 1973, CD in 2006

Songs: Sheik

CD: To A New Earth EP

Player: Kill Paris (Corey Baker)

CD Info: OWLSA, LLC (label), B00U44Z0EE, released in 2013

Songs: Slap Me

CD: The Uplift Mofo Party Plan

Player: Red Hot Chili Peppers (song original written by Bob Dylan)

CD Info: EMI (Label), B000002UD8, released in 1988

Songs: Subterranean Homesick Blues

CD: Ya-Ka-May

Player: Galactic

CD Info: Anti (Label), B0030OJPA4, released in 2010

Songs: Cineramascope

CD: Pendulum

Player: Creedence Clearwater Revival (John Fogerty)

CD Info: Fantasy (label), B001AKTZOG, released in 1970, reissue 2008

Songs: Have You Ever Seen the Rain?

CD: Electric Slave

Player: Black Joe Lewis

CD Info: Vagrant (Label), B00DOQK1TM, released in 2013 Songs: Come To My Party

CD: Lonerism

Player: Tame Impala (Kevin Parker)

CD Info: Interscope (Label), B008JFC6F0, released in 2012

Songs: Elephant

CD: Carnival Electricos

Player: Galactic

CD Info: ANTI Records (label), B006GK2WRM, released in 2012

Songs: Hey Na Na

CD: The London Souls

Player: The London Souls (Tash Neal, Chris St. Hilaire, Kiyoshi Matsuyama)

CD Info: Soul on 10 Records (label), B0053TWVUA, released in 2011

Songs: I Think I Like It

CD: The Best of Blind Blake

Player: Blind Blake

CD Info: Yazoo (Label), B00004Y9XE, released in 2000

Songs: Dry Bone Shuffle

CD: The White Album (Disc 2)

Player: The Beatles (Paul McCartney, John Lennon, George Harrison, Ringo Starr)

CD Info: Capitol (Label), B0025KVLU6, released in 1968, reissue 2009

Songs: Birthday

CD: The Chess Box: Howlin Wolf (Disc 2)

Player: Howlin’ Wolf

CD Info: Geffen (Label), B0011U39BK, released in 1991

Songs: Smokestack Lightnin’

CD: 100 Vocal & Jazz Classics – Vol. 16 (1945-1947)

Player: Dizzy Gillespie All Star Quintet

CD Info: Stardust Records (Label), B005MW5IUI, released in 2011

Songs: Salt Peanuts

CD: El Camino

Player: The Black Keys (Dan Auerbach and Patrick Carney)

CD Info: Nonesuch (label), B005URRCUY, released in 2011

Songs: Run Right Back

CD: Guns of Gold

Player: The Fumes

CD Info: Silent Partner (Label), B000GAL2P6, released in 2006

Songs: Grocery Store

CD: For the Whole World to See

Player: Death

CD Info: Drag City (label), B001NY71F4, released in 2009

Songs: Rock-N-Roll Victim

